# Mycobacterium abscessus HelR interacts with RNA Polymerase to confer intrinsic rifamycin resistance

**DOI:** 10.1101/2021.05.10.443476

**Authors:** Kelley R. Hurst-Hess, Aavrati Saxena, Pallavi Ghosh

## Abstract

Rifampicin (RIF) constitutes the frontline therapy against *M. tuberculosis* as well as most slow-growing non-tuberculous mycobacteria (NTM). However, RIF is completely ineffective against *M. abscessus* despite the absence of mutations in the rifampicin resistance determining region of *Mab_rpoB*. This has been attributed to the presence of an ADP-ribosyltransferase (Arr) activity that inactivates RIF. Rifabutin (RBT), a close analogue of RIF, has recently been shown to be effective against *M. abscessus in vitro* and in a mouse model and comprises a promising therapeutic against *M. abscessus* infections. Using RNA sequencing we show that exposure of *M. abscessus* to sublethal doses of RIF and RBT results in ∼25-fold upregulation of *Mab_helR* in laboratory and clinical isolates; an isogenic deletion of *Mab_helR* is hypersensitive to RIF and RBT, and over-expression of *Mab_helR* confers RIF tolerance in *M. tuberculosis* implying that *helR* constitutes a significant determinant of inducible RIF and RBT resistance. We demonstrate a preferential association of MabHelR with RNA polymerase *in vivo* in bacteria exposed to RIF and showed that purified MabHelR can rescue transcription inhibition in the presence of RIF in *in vitro* transcription assays. Furthermore, MabHelR can dissociate RNAP from RIF-stalled initiation complexes *in vitro*, a species we envisage accumulates upon RIF exposure. Lastly, we show that the tip of the PCh-loop of *Mab_helR*, in particular residues E496 and D497 that are in proximity to RIF, is critical for conferring RIF resistance without being required for RNAP dissociation from stalled complexes. This suggests that HelR may be additionally involved in displacing RIF bound to RNAP and function as an RNAP protection protein.

**Significance Statement:** Bacterial RNA polymerase is a target for the potent and broad-spectrum rifamycin group of antibiotics. Mutations within rpoB and inactivation by a diverse group of enzymes constitute the most widespread mechanisms of resistance. Herein we report an unprecedented mechanism of rifamycin resistance in *M. abscessus* mediated by MabHelR, a putative SF1 like helicase, that involves disassembly of RIF-stalled initiation complexes, likely followed by displacement of the antibiotic, leading to RNAP recycling. The mechanism is reminiscent of the role of HflX and ribosome protection proteins in resistance to ribosome targeting antibiotics and suggests that removal of stalled macromolecular complexes and their recycling comprises a widespread but underappreciated mechanism of antibiotic resistance. Rifampicin (RIF) is pivotal in the control of *M. tuberculosis* infections but ineffective against *M. abscessus*. Identification of inducible rifamycin resistance determinants in *M. abscessus* is therefore particularly crucial for informing treatment strategies and development of novel therapeutic approaches.

## Introduction

*M. abscessus* (*Mab*) is a rapidly growing non-tuberculous mycobacterium (NTM) and a prominent cause of broncho-pulmonary infections in patients with chronic lung damage, such as bronchiectasis, prior tuberculosis and cystic fibrosis (CF) (2013; Catherinot et al., 2013; Martiniano and Nick, 2015; Martiniano et al., 2014; Roux et al., 2009). *M. abscessus* also causes superficial and deep-tissue infections post-trauma and surgery (Kothavade et al., 2013; Nakanaga et al., 2011). It is considered an “incurable nightmare” due to its extremely low sensitivity to available antibiotics. (Ballarino et al., 2009; Brown et al., 1992a; Brown et al., 1992b; Brown-Elliott et al., 2012; Brown-Elliott and Wallace, 2002; Swenson et al., 1985; Wallace et al., 1991). Notably, *M. abscessus* displays high levels of intrinsic resistance to the anti-tuberculosis drugs, isoniazid and rifampicin (RIF) (Brown-Elliott et al., 2012; Park et al., 2008).The current treatment regimen of clarithromycin and amikacin in conjunction with cefoxitin, imipenem and tigecycline, for a period of several months (Floto et al., 2016; Griffith et al., 2007) is rendered ineffective by the rapid induction of resistance genes upon antibiotic exposure.

Rifampicin (RIF) is a potent and broad-spectrum antibiotic that comprises the first line of therapy against *M. tuberculosis* and several NTMs (Bittner and Preheim, 2016; Esteban et al., 2012). It is known to bind in a pocket of the β-subunit (RpoB) of RNA polymerase (RNAP) within the exit tunnel, thereby hindering the passage of nascent RNA when it is greater than 2-3 nt in length and an inhibition of global RNA synthesis. High levels of clinically acquired RIF resistance across bacterial species map within an 81-bp region of *rpoB* referred to as the <u>R</u>IF <u>R</u>esistance <u>D</u>etermining <u>R</u>egion (RRDR). Intrinsic RIF resistance in bacteria is attributed to the presence of refractory RNAPs, alteration of cell permeability and the presence of a diversity of enzymes that modify the drug (Goldstein, 2014; Tupin et al., 2010). RIF inactivation by ADP-ribosyl transferases (Arrs), RIF glycosyltransferases (Rgt), RIF monooxygenase (Rox), and RIF phosphorylases (Rph) have been well characterized in various actinomycetes (Spanogiannopoulos et al., 2014; Tupin et al., 2010). Additionally, the RNAP binding protein, RbpA, has been implicated to play a role in RIF resistance since a *Streptomyces coelicolor ΔrbpA* mutant is RIF sensitive and overexpression of RbpA results in increased RIF resistance in *M. smegmatis(Newell et al*., *2006)*. Although early data suggested that RbpA influences transcription inhibition by RIF on the *rrnDp3* promoter in *S. coelicolor* and in *M. smegmatis*, (Dey et al., 2010; Newell et al., 2006) subsequent studies by Hu et al.and Hurst-Hess et al. demonstrated that the apparent effect of RbpA on transcription inhibition by RIF was a reflection of significant increase in transcript yield without affecting the IC_50_ of RIF(Hu et al., 2012; Hurst-Hess et al., 2019). The effect of RbpA on RIF resistance is therefore indirect and does not involve a hinderance of RIF binding to mycobacterial RNAP (Hu et al., 2012; Hurst-Hess et al., 2019).

*M. abscessus* displays a high level of intrinsic RIF resistance despite lacking mutations in the RRDR of *rpoB* thereby rendering it unavailable for therapy against *M*.*abscessus* infections (Rominski et al., 2017). This has been attributed to the presence of an ADP-ribosyltransferase (*Mab_arr*) that ribosylates RIF thereby preventing binding to RpoB (Rominski et al., 2017). ADP-ribosyltransferases, while encoded by other fast-growing NTMs such as *M. smegmatis* and *M. fortuitum*, are absent in *M. tuberculosis*, and is the only known determinant of intrinsic RIF resistance in mycobacteria. Rifabutin (RBT), a spiro-piperidyl derivative of RIF typically exhibits lower MIC values for all mycobacteria and has been shown to be effective against *M. abscessus in vitro* and in a mouse model (Aziz et al., 2017; Dick et al., 2020; Ganapathy et al., 2019); RBT therefore constitutes a promising therapeutic option for *M. abscessus* infections. The mechanism underlying the differential potency remains to be determined but is likely a consequence of the higher lipophilicity of the drug (Blaschke and Skinner, 1996).

A transcriptomic analysis of *M. abscessus* exposed to sublethal doses of RIF revealed a >25-fold induction of *MAB_3189c*, a putative HelD-like helicase. HelD proteins belong to the SF1 superfamily of RNA and DNA helicases and are widespread in gram positive bacteria (Newing et al., 2020). Previously we showed that exposure of *M. smegmatis* to sublethal dose of RIF results in a similar upregulation of MSMEG_2174 (*Ms_helD*), the homologue of MAB_3189c (Hurst-Hess et al., 2019). Due to the involvement of *Ms_helD* and *MAB_3189c* in resistance to rifamycins, we rename these *Ms_helR* and *Mab_helR* respectively. In the present study we demonstrate that a targeted deletion of *Mab_helR* results in hypersensitivity to both RIF and RBT and investigate the mechanism of resistance. We show that MabHelR association with RNA polymerase *in vitro* rescues RIF mediated transcription inhibition and displaces RIF-stalled RNAP from bound promoter DNA. However, PCh-loop mutants of MabHelR that are proficient for dissociating stalled RNAP complexes from DNA are incapable of complementing RIF resistance of ΔMab_*helR* suggesting a potentially new mechanism of RIF resistance in which the PCh-loop of MabHelR displaces RIF bound to RNAP.

## RESUTS

### *Mab_helR* expression is induced by sublethal concentrations of RIF and RBT

In a preliminary study we used RNA sequencing (RNAseq) to determine the genome-wide changes in gene expression in *Mab* ATCC 19977 upon exposure to sublethal doses of RIF (16ug/ml for 30 mins). We observed that six genes were upregulated > 4-fold (*p*_*adj*_ <0.01) (Figure 1a). Of these *Mab_arr* and MAB_3189c showed the greatest changes in expression (>25-fold) which were confirmed using real time PCR (Figure 1b). MAB_3189c is a homologue of HelD proteins that belong to the SF1 family of RNA and DNA helicases, and is referred here as *helR*. Further, we showed that *Mab_helR* and *Mab_arr* were similarly upregulated upon exposure to sublethal doses of RBT, a RIF analogue with *in vitro* activity against *M. abscessus* (Figure 1b). In order to determine the clinical relevance of *helR* in rifamycin resistance, we followed the RIF/RBT resistance and expression of *helR* in 8 sequenced clinical isolates of *M. abscessus* obtained from the from the Wadsworth Center Reference Laboratory. The eight isolates included all three subspecies of *M. abscessus*-*M. abscessus abscessus, M. abscessus masiliense* and *M. abscessus bollettii* and share 96.6-99.8% average nucleotide identity (ANI) (Figure 1c); all strains displayed rifamycin tolerance similar to that of *Mab* ATCC (Table S1). As seen in Figure1d, exposure to sublethal concentrations of RIF resulted in upregulation of both *Mab_arr* and *Mab_helR* in all eight clinical isolates of *M. abscessus*. The upregulation of *Mab_arr* upon RIF exposure is consistent with previous studies which show that Arr is a major determinant of RIF resistance in *M. abscessus* (Rominski et al., 2017). A similar upregulation of *Mab_helR*, in laboratory and clinical strains, is therefore suggestive of a significant role of MabHelR in rifamycin resistance.

**Figure 1.**
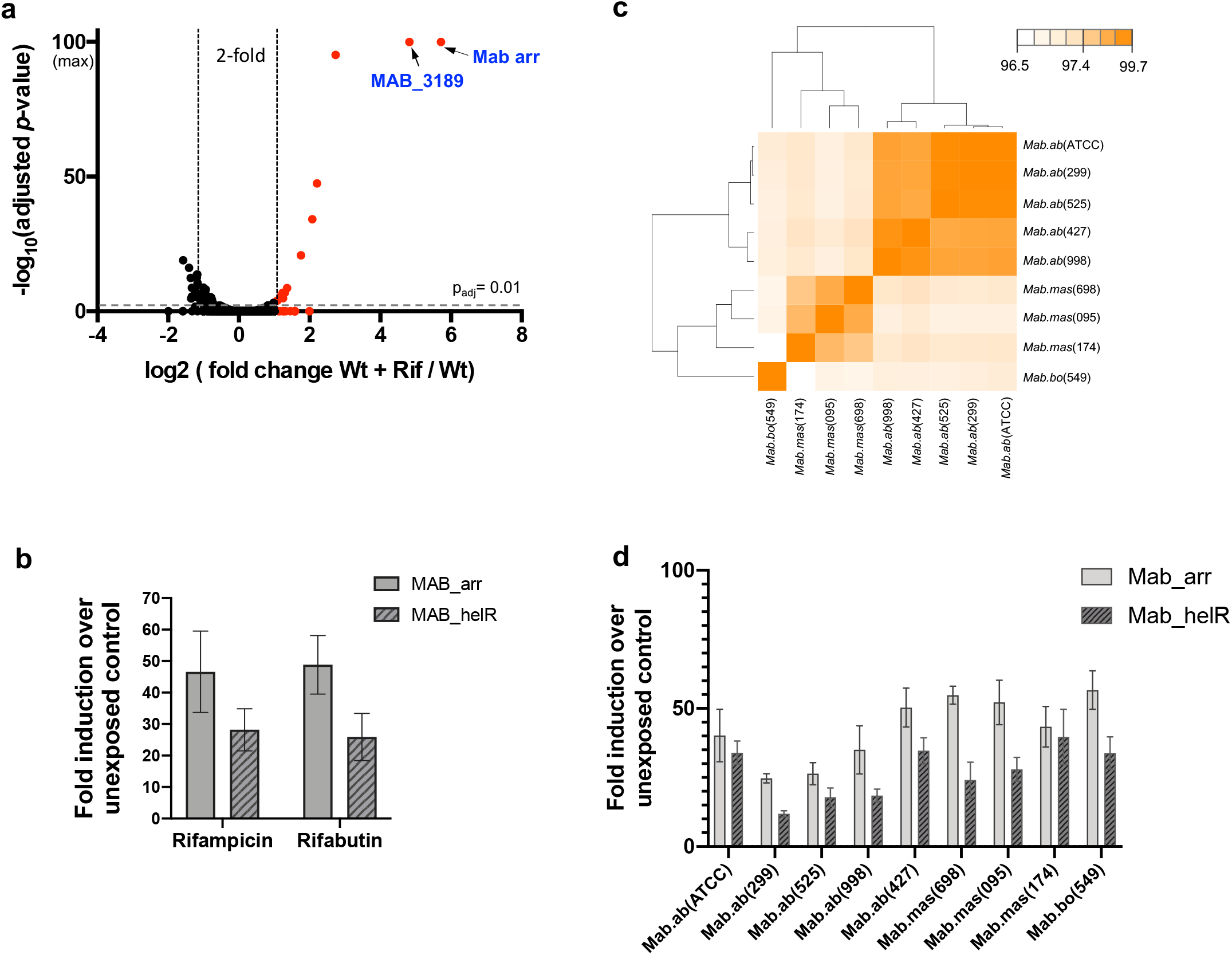
Exposure of *M. abscessus* ATCC and clinical strains to rifamycins results in induction of MAB_3189c/ *helR* expression. **a**) Volcano plot of differentially expressed genes in *M. abscessus* ATCC19977 upon exposure to RIF (16μg/mL for 30 mins). *Mab_arr* and *MAB_3189* (*helR*) are the two most highly induced genes and are indicated. **b**) Wild type *M. abscessus* was grown to A_600_ of 0.7 and exposed to either 8μg/mL of RIF or 0.5 μg/mL of RBT for 30 mins. The amount of *Mab_arr* and *Mab_helR* transcript was determined by qPCR and plotted as fold induction over an unexposed control. Data represents mean ± SD, n=3. *SigA* was used as an internal normalization control. **c**) Average nucleotide identities between eight clinical strains of *M. abscessus* obtained using FastANI version 1.32. is shown. **d**) Expression of *Mab_arr* and *Mab_helR* transcript in eight clinical strains of *M. abscessus* upon exposure to 8μg/mL of RIF determined by qPCR and plotted as fold induction over an unexposed control. Data represents mean ± SD, n=3. *SigA* was used as an internal normalization control.

### Deletion of *Mab_helR* results in hypersensitivity to RIF and RBT

An isogenic deletion of *Mab_helR* in MabATCC was created using phage recombineering (Hurst-Hess et al., 2017). As seen in Figure 2a, a *ΔMab_helR* strain was hypersensitive to RIF and RBT. Expression of *Mab_helR* driven by a native promoter from a chromosomally integrated copy in the *ΔMab_helR* strain restored sensitivity of the mutant to wild-type levels (Figure 2a, Table 1). The growth of *ΔMab_helR* in media lacking RIF remained unchanged (Figure S1a). Moreover, a double mutant of *ΔMab_helR* / *ΔMab_arr* was more sensitive than each single mutant alone suggesting that *Mab_helR* and *Mab_arr* comprise independent and additive resistance mechanisms (Figure 2b, Table 1). Additionally, the expression of *Mab_arr* remained unchanged in a *ΔMab_helR* background, confirming that the observed hypersensitivity of *ΔMab_helR* to rifamycins is not a consequence of a decrease in expression of *Mab_arr* in the *ΔMab_helR* strain (Figure 2c).

**Table 1.** Survival of wild type *M. abscessus* ATCC19977, *ΔHelR, ΔMab_arr, ΔMab_helR/ ΔMab_arr, ΔhelR +*pMH94-*helR* in a 2-fold dilution series of RIF and RBT in Middlebrook 7H9/OADC medium. The minimum concentration of antibiotic required to inhibit 99 % of growth after 72 hours is shown. Data is representative of 3 replicates.

| Strain | Minimum Inhibitory Conc. (µg/mL) |  |
| --- | --- | --- |
|  | RIFAMPICIN | RIFABUTIN |
| WT *Mab* | 64 | 2 |
| Δ*Mab_helR* | 8 | 0.5 |
| Δ*Mab_arr* | 1 | 0.125 |
| Δ*Mab_helR*/Δ*Mab_arr* | 0.125 | 0.0039 |
| Δ*Mab_helR* + pMH*Mab-helR* | 32-64 | 2 |

**Figure 2.**
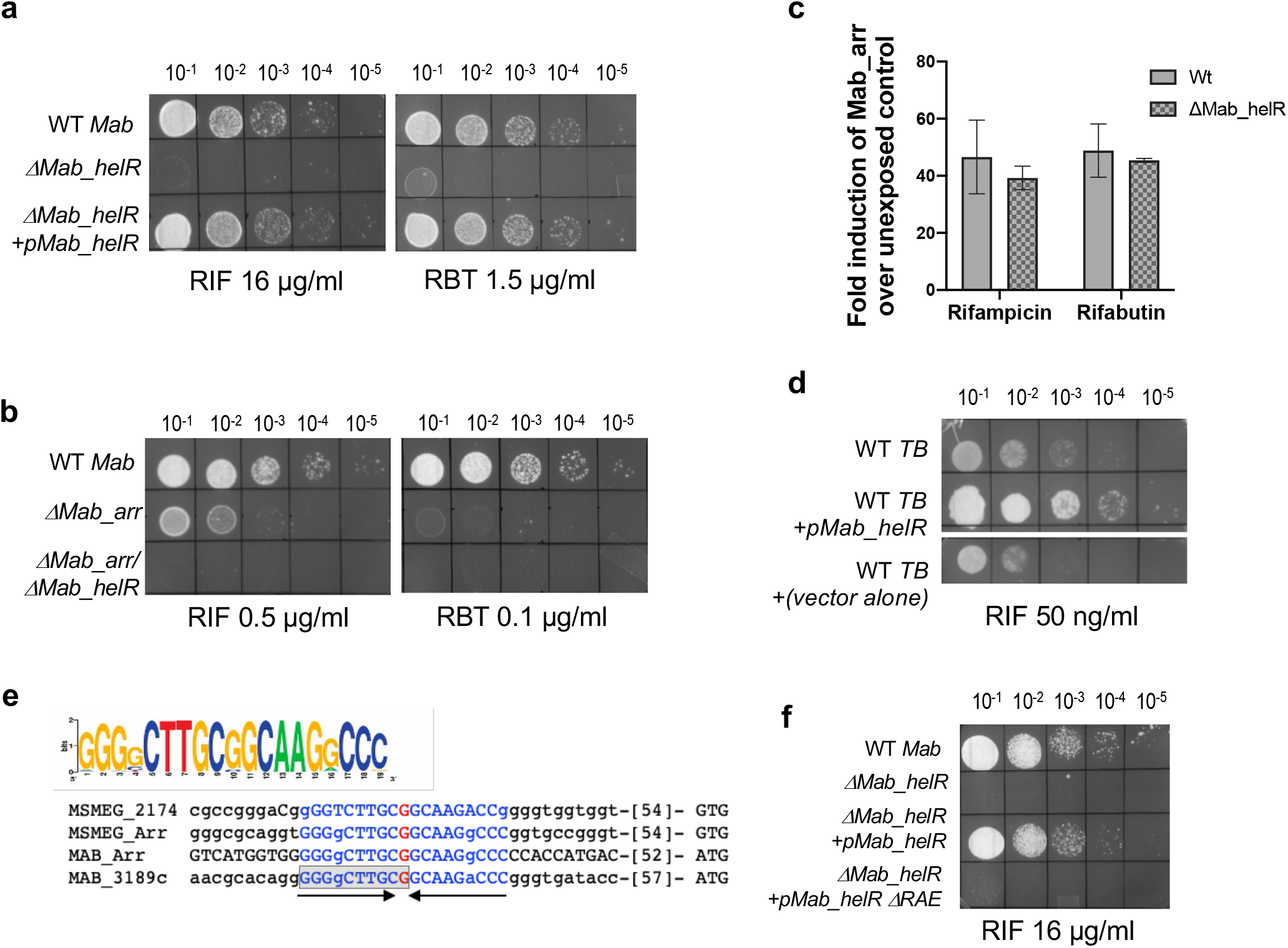
Deletion of *Mab_helR* results in RIF and RBT hypersensitivity in *M. abscessus* and heterologous expression of *HelR* in *M. tuberculosis* increases RIF tolerance. **a-b)** Growth of ten-fold serial dilutions of *M. abscessus* ATCC 19977, *ΔMab_helR, ΔMab_arr,ΔMab_helR/ ΔMab_arr*, and the indicated complementing strains on Middlebrook 7H10 OADC containing indicated concentrations of RIF and RBT. Data is representative of >3 independent experiments. **c**) Real time PCR showing the induction over an unexposed control of *Mab_arr* transcript in wild type *M. abscessus* and the *ΔhelR* strains exposed to either 8μg/mL of RIF or 0.5 μg/mL of RBT. Data represents mean ± SD, n=3. *SigA* was used as an internal normalization control. **d**) Growth of ten-fold serial dilutions of wild-type *M. tuberculosis mc*^*2*^*7000, mc*^*2*^*7000* containing a chromosomally integrated copy of *helR* and *mc*^*2*^*7000* containing a chromosomally integrated copy of the empty vector (control) on Middlebrook 7H10 OADC/pan 50ng/mL of RIF. Data is representative of 2 independent replicates. **e**) Sequence logo of the19bp inverted repeat sequence (RAE) and alignment of upstream sequences of *Mab_helR* (MAB_3189), *Mab_arr, Ms_helR* (MSMEG_2174) and *Ms_arr*. Region of RAE deleted is shaded in grey. **f**) Growth of ten-fold serial dilutions of *M. abscessus* ATCC 19977, *ΔMab_helR* and complementing strains containing either *helR* expressed from a native promoter or *helR* expressed from a promoter lacking one half of RAE on Middlebrook 7H11 OADC containing RIF (16μg/mL).

*M. tuberculosis* H37Rv is highly susceptible to RIF and lacks homologues of *Mab_arr* and *Mab_helR*. We expressed *Mab_helR* in the RIF sensitive *M. tuberculosis mc*^*2*^*7000*, a severely attenuated derivative of *M. tuberculosis* H37Rv (Ojha et al., 2008). As seen in Figure 2d, heterologous expression of *Mab_helR* resulted in increased RIF tolerance of *M. tuberculosis mc*^*2*^*7000*. Furthermore, *M. smegmatis ΔMs_helR* is also RIF sensitive and expression of *Mab_helR* in a *ΔMs_helR* strain restored its antibiotic tolerance to wild-type levels thereby suggesting a conserved function of *Mab_helR* and *Ms_helR* (Figure S1b)(Hurst-Hess et al., 2019). These results together establish the role of *HelR* as an additional mechanism of RIF resistance in mycobacteria and demonstrate that *Mab_helR* and *Mab_arr* constitute inducible mechanisms of basal RBT tolerance in *M. abscessus*.

An analysis of upstream sequences of *Mab_helR* revealed the presence of the RIF associated element (RAE), a highly conserved 19-bp inverted repeat sequence, previously identified upstream of genes encoding RIF-inactivating enzymes (glycosylation, ADP ribosylation and monooxygenation) and helicases from divergent actinomycetes (Spanogiannopoulos et al., 2014) (Figure 2e). The conserved RAEs were found upstream of only *Mab_arr* and *Mab_helR* within the *M. abscessus* genome underscoring the importance of *helR* in rifamycin resistance and suggesting a co-regulation of *Mab_arr* and *Mab_helR* by a conserved mechanism. Deletion of one half of the conserved RAE (Figure 2e, shaded in grey) results in the inability to restore RIF sensitivity of the *ΔhelR* strain (Figure 2f) confirming a role for RAE in the induction of RIF resistance.

### MabHelR interacts with RNA polymerase *in vivo and* rescues RIF inhibition of RNAP in *in vitro* transcription assays

Previous studies on a putative RAE-associated helicase from Streptomyces WAC4747 demonstrated its inability to inactivate RIF and suggested a mechanism of action other than RIF modification (Spanogiannopoulos et al., 2014). HelD was first detected as a major copurifying protein in preparations of RNA polymerase from *B. subtilis*, and has since been established as a direct binding partner of *B. subtilis* RNA polymerase involved in RNAP cycling (Newing et al., 2020; Pei et al., 2020; Wiedermannova et al., 2014). Recently, the cryoEM structure of *M. smegmatis* HelR in complex with RNA polymerase also suggested a role of MsHelR in dissociating stalled elongating RNAP complexes from the associated nucleic acids (Kouba et al., 2020). In order to determine if MabHelR similarly associates with RNAP, we constructed a MabATCC strain containing a 10X his-tag at the native *rpoA* locus. MabRNAP_his_ was purified from cells exposed to sublethal quantities of RIF as well as from untreated cells. A band corresponding in size to purified MabHelR was found to be ∼5-fold enriched in RNAP preparations from RIF-exposed cells and was identified as MabHelR using LC-MS mass spectrometry (Figure 3a). Negligible amounts of MabHelR were detectable in RNAP purified from exponentially grown cells in the absence of antibiotic (Figure 3a).

**Figure 3.**
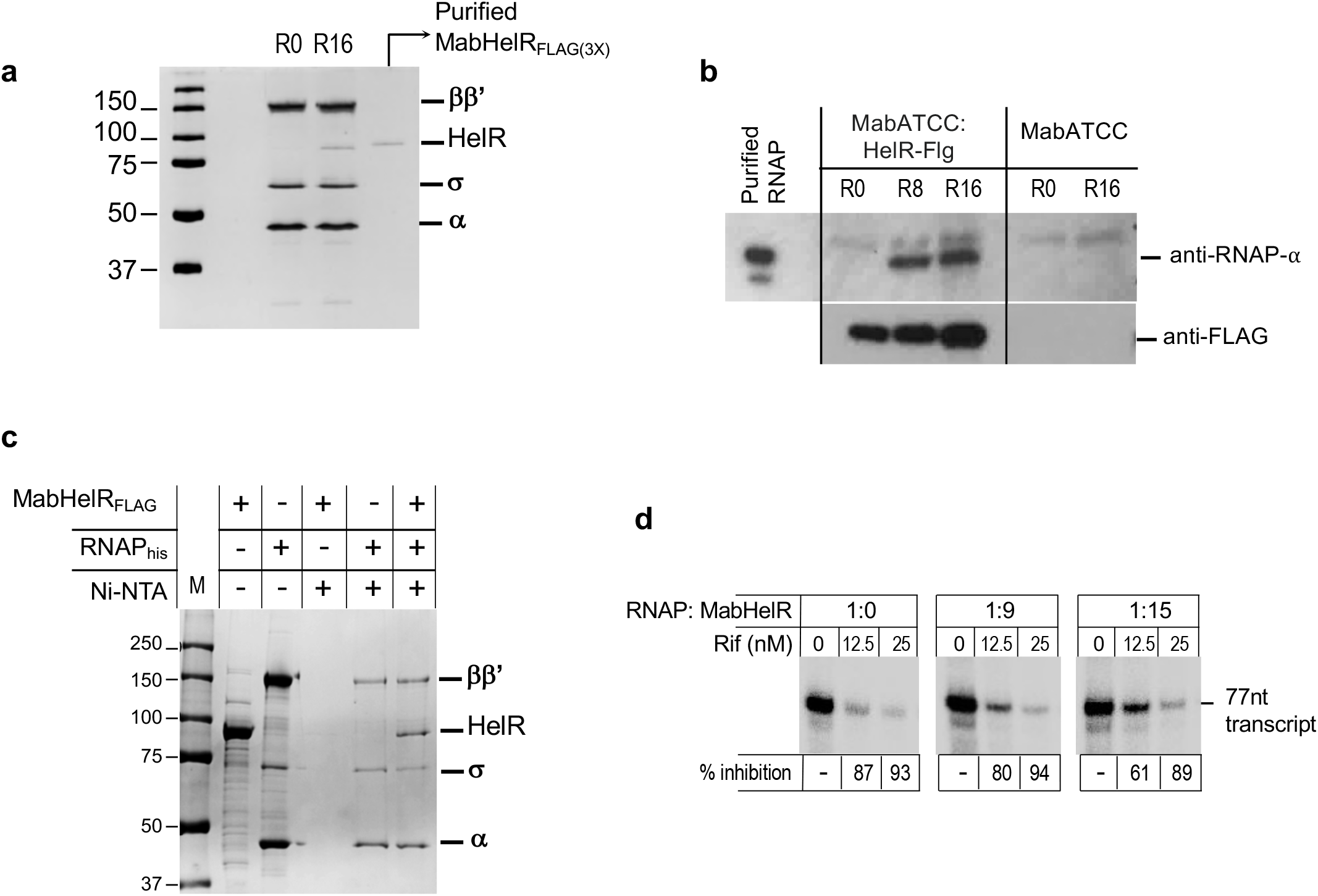
MabHelR associates with RNAP *in vivo* and *in vitro* and rescues inhibition of *in vitro* transcription in the presence of RIF. **a**) Silver stained gel showing RNAP purified from untreated and RIF treated (16μg/mL of RIF for 30 mins) *M. abscessus* strain in which *rpoA* was C-terminally tagged with the 10X-his epitope at its native chromosomal location. Purified MabHelR is used as a control. RNAP purified from RIF treated samples was loaded on a preparative SDS PAGE, Coomassie stained and a band corresponding in size to MabHelR was excised and analyzed by LC MS/MS. **b**) An *M. abscessus* strain in which *helR* was C-terminally tagged with the 3X-FLAG epitope at its native chromosomal location was either untreated or treated with 8-16μg/mL of RIF for 30 mins. Wild-type *M. abscessus* with an untagged *helR* gene was used as a control. Samples were normalized by wet cell weight and FLAG tagged proteins were purified using anti-FLAG M2 beads. Samples were analyzed by immunoblotting using anti-FLAG and anti-RpoA antibodies. Purified RNAP was used as a marker. Data is representative of >3 biological replicates. **c**) Coomassie stained gel showing elution of proteins using nickel affinity chromatography. Interaction assays were carried out using HelR_FLAG_ and RNAP_his_ followed by binding and elution from a Ni-NTA column. Control samples included either only RNAP_his_ or HelR_FLAG._ Purified RNAP_his_ or HelR_FLAG_ are included as markers. **d**) Multiple-round *in vitro* transcription assays were performed on the *sinP3* promoter using 75nM σ^A^-RNAP. RIF was added to indicated concentrations for 10 mins at 37°C followed by addition of a 9-or 15-fold molar excess of MabHelR. Transcription was initiated by addition of NTPs and incubated at 37°C for 30 mins. Samples were separated using denaturing PAGE (6% Urea polyacrylamide gel) and the 77nt product was visualized using a Typhoon Imager (GE Healthcare) and quantitated using the Image Quant software. The inhibition of transcription at each RIF concentration is expressed as a ratio of the activity in the absence and presence of RIF.

Next, MabHelR was purified from a MabATCC strain containing a 3X FLAG-tag at the native *helR* locus exposed to sublethal quantities of RIF, as well as from RIF untreated cells. Western blot analysis with anti-FLAG antibody showed the presence of significant quantities of cellular HelR_FLAG_ in antibiotic-untreated *M. abscessus* which increased in RIF treated cells (Figure 3b). However, interaction of MabHelR with MabRNAP was detectable using anti-RNAP-α antibody predominantly in RIF-treated cells (Figure 3b). While this does not rule out interaction of HelR with RNAP in exponentially growing bacteria, it demonstrates an enhanced association with RNAP in the presence of RIF.

To validate that purified MabHelR interacts directly with RNAP, we performed *in vitro* protein-protein interaction assays using purified HelR_FLAG_ and his-tagged RNAP. As seen in Figure 3c, coelution of HelR_FLAG_ with RNAP_his_ from a Ni-NTA column was observed only in the presence of RNAP; in the absence of RNAP, MabHelR_FLAG_ was not retained on a Ni-NTA column implying that MabHelR interacts directly with RNAP.

Finally, since a deletion of *HelR* results in hypersensitivity to rifamycins, we wished to study the effect of MabHelR on RIF mediated transcription inhibition. Multiple round *in vitro* transcription was performed using *M. tuberculosis* RNAP that naturally lacks HelR, and the well-characterized *sinP3* promoter (Jacques et al., 2006). A 77nt run-off transcript was observed that diminished in the presence of increasing concentrations of RIF. Addition of a 9-fold or 15-fold molar excess of MabHelR resulted in increased transcript formation at each of the RIF concentrations tested as compared to control reactions lacking HelR suggesting that MabHelR could rescue transcription inhibition of RNAP by RIF (Figure 3d).

### The ATPase activity of MabHelR is required for RIF resistance

HelD-like proteins display a four-domain structure containing an N-terminal domain, a superfamily 1 (SF1) 1A domain that is split into a 1A-1 and a 1A-2 at the primary amino-acid sequence level by a HelD specific domain, and a C-terminal 2A domain. Similar to *M. smegmatis* HelR, the four conserved structural domains could be discerned in MabHelR (Figure 4a, Figure S2). Domains 1A and 2A together form a conserved Rossman fold with seven conserved motifs, including Walker A and Walker B that are required for NTP binding and hydrolysis (Caruthers and McKay, 2002). Purified MabHelR displayed significant ATPase activity that is independent of RNAP (Figure 4b) and was consistent with the behavior of *B. subtillus* HelD (Wiedermannova et al., 2014). In order to determine if the ATPase activity of MabHelR is required for its ability to confer RIF resistance, we generated two variants of MabHelR that contained mutations in either the Walker B motifs (MabHelR-mWalkerB) or both Walker A and B motifs (MabHelR-mWalkerAB). Both mutant proteins were defective in ATP hydrolysis *in vitro* (Figure 4b). Further, MabHelR-mWalkerB and MabHelR-mWalkerAB were unable to restore rifamycin sensitivity of *ΔMab_helR in vivo* despite retaining the ability to bind RNAP (Figure 4c, Figure S3a) implying that ATP hydrolysis is necessary for HelR mediated RIF resistance. Control experiments showed that upon RIF exposure MabHelR-mWalkerB and MabHelR-mWalkerAB were expressed from the integrated chromosomal copy at levels similar to that of WT HelR (Figure S3b).

**Figure 4.**
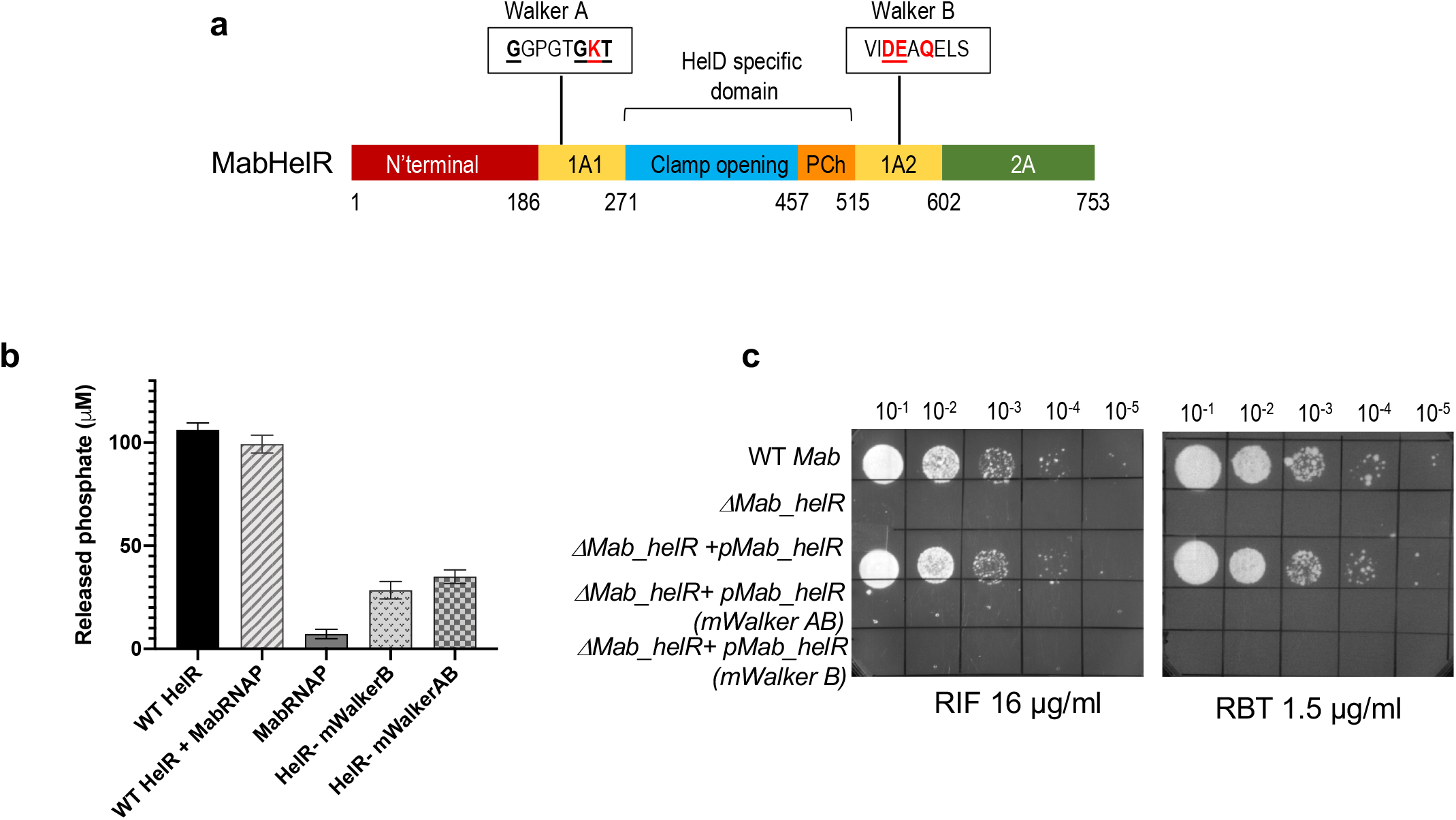
MabHelR is an ATPase and ATP hydrolysis is required for conferring RIF/RBT resistance. **a**) Domain organization of MabHelR showing the location of Walker A and Walker B motifs within domains 1A1 and 1A2. The strictly conserved residues in each motif are underlined and the residues mutated to alanine in the study are in red. **b**) ATPase activity of Wt MabHelR is shown in the presence and absence of MabRNAP. MabHelR mutations in either Walker B alone (mWalkerB) or Walker A and B (mWalkerAB) are defective in ATP hydrolysis. Data represents mean ± SD, n=3 **c**) Growth of ten-fold serial dilutions of *M. abscessus* ATCC 19977, *ΔMab_helR* and *ΔMab_helR* complemented with either *WT helR, helR* (mWalkerB) or *helR* (mWalkerAB) on Middlebrook 7H10-OADC containing RIF (16 μg/mL) or RBT(1.5 μg/mL). Data is representative of at least 3 independent experiments.

### The PCh loop of MabHelR is required for RIF resistance but not for dissociation of RIF-stalled RNAP complexes from DNA

The cryoEM structure of MsHelR revealed a crescent shaped structure; one end of the crescent formed by the N-terminal domain inserts into the secondary channel, while the other end formed by the clamp opening (CO) domain and the primary channel (PCh) loop, inserts into the primary channel (Kouba et al., 2020). Upon binding RNAP, the PCh loop folds into a helix α16 and a helix α17 that pack against the β’ bridge helix (BH) and is inserted into the active site (AS) cavity. A model of MabHelR was generated using Phyre v2.0 and superimposed on the structure of MsHelR-RNAP complex (PDB ID 6YYS). Superimposition of the crystal structure *of M. smegmatis* RNAP initiation complex bound to RIF (PDB ID 6CCV) was used to locate RIF on the RNAP-HelR complex (Figure 5a). The tip of the PCh-loop consists of four acidic residues, MabHelR/494-DDED-497. Of these, residues MabHelR/Asp494 and Asp495 are in proximity to MgA and AS conserved motif β’ – NADFDGD, while Glu496 and Asp497 are in close proximity (within 3.5 Å) to RIF. Due to the proximity of the PCh-loop to RIF, we sought to investigate the role of the PCh-loop in the ability of MabHelR to confer RIF resistance. For this purpose we generated three mutants of MabHelR; ΔPCh1: deletion in aa 467-515, ΔPCh2: deletion in aa 486-505, and mPCh: E496A/D496A. As seen in Figure 5b, deletion of either the entire PCh loop α16 and α17 (ΔPCh1) or the intervening region between the helices (ΔPCh2) results in an inability to restore rifamycin sensitivity of *ΔMab_helR in vivo*, despite retaining the ability to bind RNAP *in vitro* (Figure S3a). Importantly, alanine substitution of Glu496 and Asp497 is sufficient to result in a marked defect in the ability of the MabHelR-mPCh to confer RIF resistance (Figure 5b). Control experiments showed that RIF inducible expression of MabHelR ΔPCh1,ΔPCh2 and mPCh in the complementing strains were similar to that of WT MabHelR (Figure S3b).

**Figure 5.**
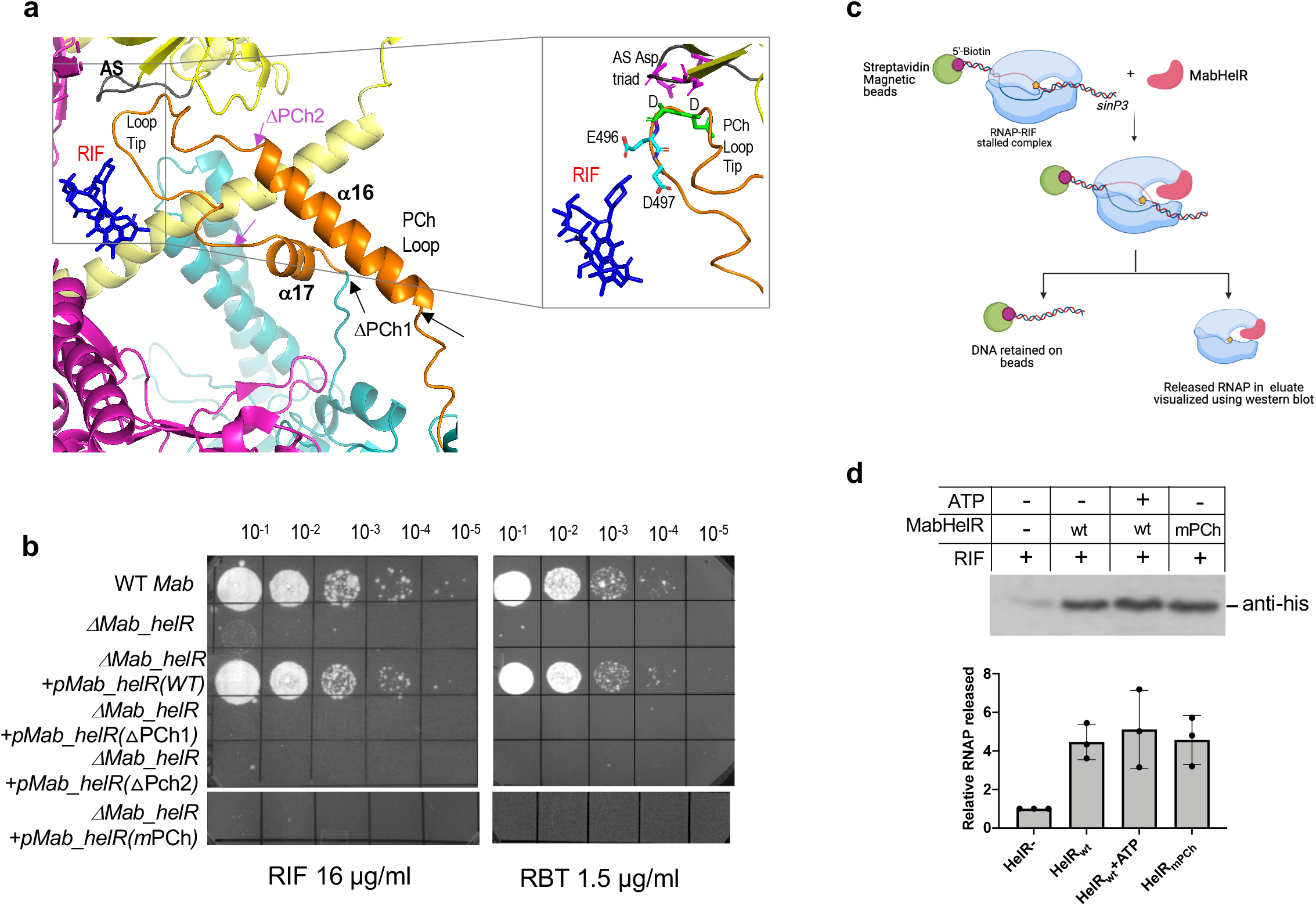
PCh loop of MabHelR is essential for conferring RIF/RBT resistance. **a**) Location of the MabHelR PCh loop (orange) containing helices α16 and α17relative to the active site (AS) residues (gray) in β’ (yellow) and RIF (blue) is shown. Structure of MabHelR was modeled using Phyre and superimposed on the structure of MsHelR bound to RNAP (PDB ID 6YYS) in PyMOL v2.3. 0. The location of RIF was determined from the structure of RIF bound *M. smegmatis* RNAP (PDB ID 6CCV). Black arrows show the regions of MabHelR PCh loop that are deleted in MabHelRΔPCh1 and magenta arrows show the deleted regions in MabHelR ΔPCh2. Enlarged box below shows the location of the 494-DDED-497 motif at the tip of the PCh loop and the proximity of E496/ D497 to RIF. **b**) Growth of ten-fold serial dilutions of *M. abscessus* ATCC 19977, *ΔMab_helR* and *ΔMab_helR* complemented with either *WT helR, helR* (ΔPCh1) *helR* (ΔPCh2) and *helR* (mPCh) on Middlebrook 7H10-OADC containing RIF (16 μg/mL) or RBT(1.5 μg/mL). Data is representative of at least 3 independent experiments. ΔPCh1 corresponds to deletion of residues 467-515, ΔPCh2: deletion of residues 486-505 and mPCh corresponds to point mutations E496A/D496A. **c**) Schematic representation of experimental design. RNAP stalling on 5’-bio *sinP3* DNA was carried out in the presence of RIF and ATP, the +1 nucleotide. The stalled complexes were tethered on streptavidin magnetic beads. Upon addition of MabHelR, the dissociated RNAP is expected to be present in the eluate and RNAP bound to DNA remains tethered on the beads. **d**) Western blot of samples eluted from streptavidin magnetic beads tethered to DNA with RIF trapped RNAP complexes using anti-his antibodies that recognize his-tagged RpoA. Western blots are quantitated using ImageJ. Data represents mean ± SD, n=3.

Previous studies have shown that the BsuHelD and MsHelR can release stalled RNAP elongation complexes from bound DNA. Binding of RIF to the β subunit of RNAP within the DNA/RNA channel sterically blocks the passage of the RNA transcript at the 5′ end when the transcript becomes either 2 or 3 nt in length, without affecting substrate binding, catalytic activity or translocation (Campbell et al., 2001). RNAP inhibited in the presence of RIF conceivably constitute stalled initiation complexes that could be targets of HelR. In order to determine if MabHelR can dissociate RIF stalled RNAP complexes from bound promoter DNA, we assembled initiation complexes on a 5’-biotinylated *sinP3* template stalled in the presence of RIF and the initiating nucleotide, ATP. The DNA-RNAP complex was tethered on streptavidin agarose beads and the dissociation of stalled RNAP from promoter DNA was studied as a function of MabHelR addition using western blot analysis with anti-his antibodies that recognize RpoA_his_ (Figure 5c). MabHelR was able to dissociate stalled RIF-RNAP complexes from DNA independent of the presence of ATP which is consistent with previous experiments with stalled *M. smegmatis* elongation complexes (Figure 5d) (Kouba et al., 2020). Additionally, the mPCh mutant was also proficient in disassembly of stalled RIF-RNAP complexes despite its inability to complement the RIF sensitivity of the *ΔMab_helR* mutant, suggesting that HelR mediated RIF resistance requires a step in addition to displacement of stalled initiation complexes, likely the active displacement of RIF mediated by interactions with the PCh-loop.

## Discussion

Enzymatic inactivation of RIF by the rifampicin ADP-ribosyl transferase (*arr)* is known to be the primary determinant that contributes to the high levels of innate RIF resistance of *M. abscessus*. In the present study we identify a new and significant determinant of inducible RIF/RBT resistance in *M. abscessus*, MabHelR, that functions via an *arr* independent pathway. We demonstrate that unlike BsuHelD, which is seen to associate in stoichiometric amounts with RNAP isolated from exponentially growing bacteria (Wiedermannova et al., 2014), MabHelR preferentially associates with RNAP isolated from RIF treated samples (Figure 3 a-b). The increased association of MabHelR with RNAP from RIF treated bacteria could be a mere reflection of increased transcription of *Mab_helR* upon RIF exposure. However, we note that the amount of cytosolic MabHelR protein does not show an increase proportionate to *Mab_helR* transcription upon drug addition (Figure 3b). The inability to detect an interaction of HelR with RNAP in exponentially growing bacteria despite the presence of significant quantities of cytosolic HelR suggests that RIF bound RNAP constitutes a prime target of MabHelR.

The cryo-EM structure of MsHelR bound to RNAP reveals that the N-terminal domain is inserted into the secondary channel of RNAP in a way that interferes with the nucleotide addition cycle, and the primary channel is occupied by the PCh-loop and the CO domain (Kouba et al., 2020). The PCh-loop penetrates into the AS and leads to a repulsion of the dsDNA and RNA/DNA hybrid at the AS while the CO domain binds the β’-clamp thereby splaying open the RNAP primary channel. Together, these interactions lead to spillage of nucleic acid contents from a transcription complex. The interaction of HelR with RNAP is conceivably inhibitory for transcription and is thought to be required for dissociation of RNAP complexes that are stalled either during elongation or termination. We hypothesized this HelR-RNAP interaction can be extended to the RNAP holoenzyme stalled during initiation in the presence of transcription inhibitors like RIF (Figure 6, complex A), a scenario compatible with previous observations of Kouba *et al*. that RNAP, MsHelR and σ^A^ can coexist within one complex (Kouba et al., 2020). The ability of MabHelR to dissociate RNAP trapped in the presence of RIF on the *sinP3* promoter DNA, as well as alleviation of transcription inhibition by RIF in multiple round *in vitro* transcription assays provide support for this model (Figures 3d and 5c,d). Since the RIF-stalled complexes constitute significant roadblocks for the transcription-translation machinery and replication, the clearance of these obstacles is imperative for bacterial survival offering a possible explanation for the RIF hypersensitivity of the *ΔhelR* mutant (Figure 2).

**Figure 6.**
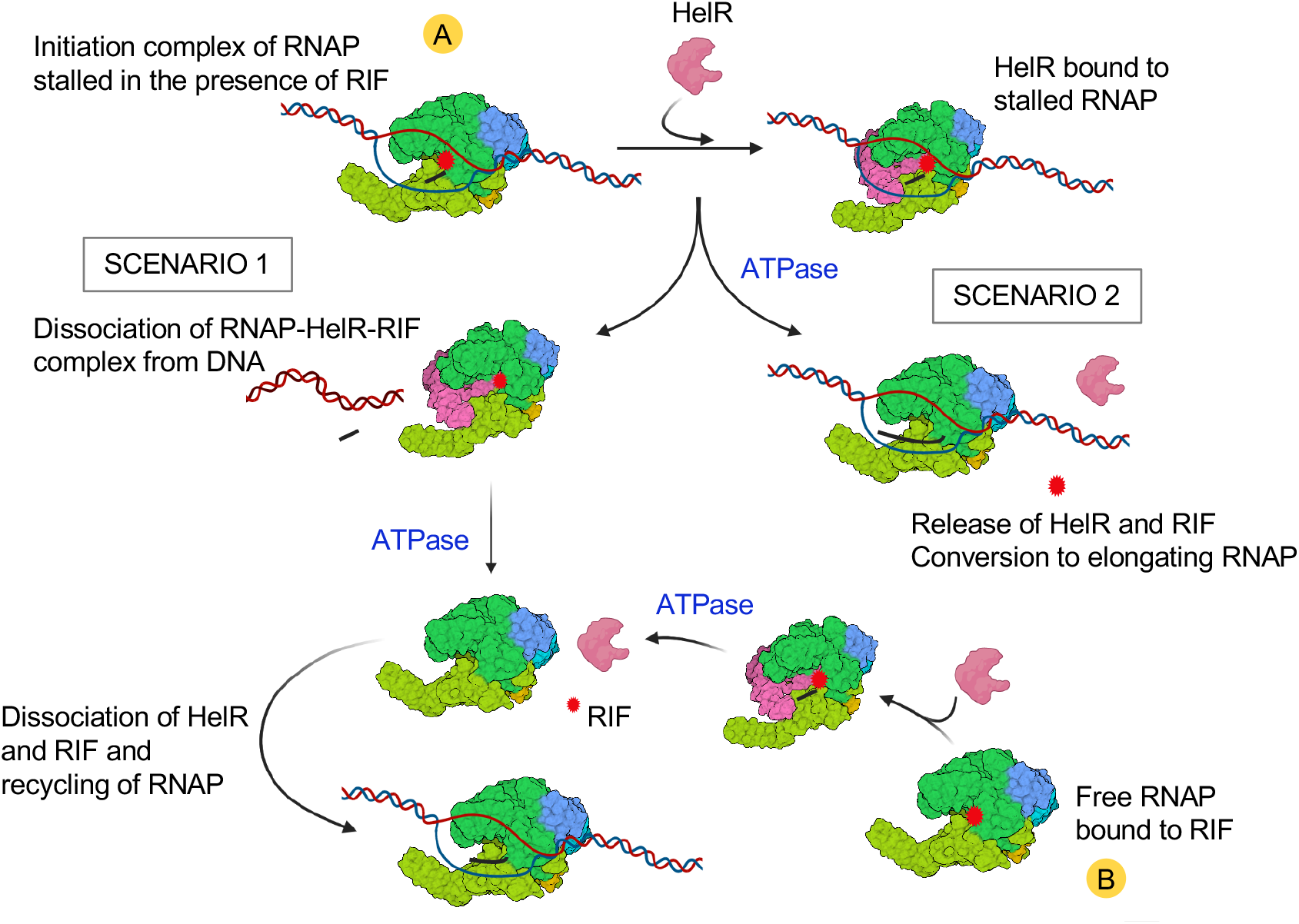
Model of HelR mediated RIF resistance. Two species of RNAP that can serve as substrates for HelR and are labeled as (A) RNAP stalled during initiation in the presence of RIF and (B) free RNAP. Possible pathways for HelR action leading to RNAP recycling are illustrated.

Interestingly, data in Figure 5d demonstrate that MabHelR can dissociate RIF-stalled RNAP in an NTP independent manner and is consistent with previous *in vitro* studies that describe an NTP independent disassembly of stalled elongation complexes by MsHelR (Kouba et al., 2020). Nevertheless, MabHelR mutants defective in ATP hydrolysis are also defective in conferring rifamycin resistance implying that ATP hydrolysis must be required for RIF resistance in a step other than disassembly of the RNAP complex (Figure 4). Additionally, removal of the tip of the PCh-loop (MabHelR486-505), specifically mutations in MabHelRE496 and D497 that are in close proximity to RIF (within 3.6 Å), abrogates the ability of MabHelR to restore rifamycin resistance of a *Δ Mab_helR* mutant but does not affect its ability to dissociate stalled RNAP (Figure 5d). Taken together these data suggest that disruption of stalled RNAP by MabHelR is insufficient for HelR mediated RIF resistance. We therefore envision that MabHelR may actively displace RIF bound to RNAP following dissociation of the stalled initiation complex (Figure 6, Scenario 1) and is corroborated by studies on the *Streptomyces venezuelae* HelR which demonstrate displacement of RIF from RNAP using a RIF photoprobe (Gerard Wright, personal communication). The requirement of ATPase activity of MabHelR in RIF resistance suggests that displacement of RIF and/or dissociation of HelR from RNAP may be dependent on ATP hydrolysis. An alternate pathway (Figure 6, scenario 2) could involve removal of RIF by HelR binding to RIF-trapped initiation complexes, dissociation of HelR and resumption of transcription. This model is however unlikely due to the instability of the RNAP-DNA complex induced by HelR binding. Finally, we cannot rule out that free RNAP bound to RIF could also serve as substrates for HelR, and displacement of RIF by HelR makes RNAP available for further rounds of transcription (Figure 6, complex B).

The function of HelR in RIF resistance, while unprecedented for RNAP, displays a startling resemblance to HflX and the ribosome protection proteins that confer resistance to ribosome targeting antibiotics. Recently we demonstrated that MabHflX confers resistance to macrolide/lincosamide antibiotics by dissociating ribosomes stalled in the presence of these drugs (Rudra et al., 2020). A similar mechanism of HflX mediated macrolide/lincosamide resistance has also been suggested for *Listeria monocytogenes* (Duval et al., 2018). The clearance of antibiotic stalled macromolecular complexes therefore appears to be a common but underappreciated theme of resistance mechanisms employed by bacteria. Additionally, ribosome protection proteins such as TetM/TetO involved in tetracycline resistance, and the ARE-ABCF proteins involved in resistance to antibiotics that bind the peptidyl transferase center/ nascent polypeptide exit tunnel act by displacing the ribosome bound antibiotic (Connell et al., 2003; Sharkey et al., 2016). The HelR mediated rifamycin resistance mechanism therefore appears to integrate the functions of both HflX and the ribosome protection proteins within a single protein.

The inducible expression of *Mab_arr* and *HelR* renders RIF completely ineffective as a therapeutic option against *M. abscessus* infections. RBT induced expression of *Mab_arr* and *Mab_helR* could potentially limit the efficacy of RBT treatment as well. Interestingly, the presence of the conserved 19-bp inverted repeat RAE sequence identified upstream of both *Mab_arr* and *Mab_helR* suggests the presence of a common regulator. Since a double mutant is more sensitive than a single mutant of either *Mab_arr* or *Mab_helR* alone, targeting the common regulator can potentially reduce RIF sensitivity dramatically and reclaim the use of this well tolerated antibiotic. Future characterization of RAE-dependent regulation of *Mab_arr* and *Mab_helR* will be of paramount importance as it will enable development of inhibitors that can concomitantly prevent expression of both rifamycin resistance determinants leading to an effective therapeutic regimen containing a rifamycin antibiotic.

## MATERIALS & METHODS

### Bacterial Strains, Media and Plasmids

*M. abscessus* ATCC19977 strains were grown at 37°C in Middlebrook 7H9 (DIFCO) supplemented with 0.05% Tween 20 and 10% OADC/ADC. Gene replacement mutants as well as tagged strains were constructed using recombineering and confirmed by sequencing (Hurst-Hess et al., 2017). Complementing strains were created by cloning WT or mutant *HelR* in pMH94 under the control of its native promoter followed by integration at the phage L5 *attB* site. Wt and mutant *HelR* genes were cloned in pET21a or pET21a-FLAG for overexpression in *E*.*coli*. were Deletions and point mutations in *HelR* were created using the Q5 site directed mutagenesis kit (NEB). Primers for mutant construction were designed using the NEBaseChanger as recommended. All mutants were confirmed by Sanger sequencing. All bacterial strains and plasmids are listed in Table S2.

Genomes of 9 clinical strains *M. abascessus* were sequenced using the Illumina platform and assembled using Unicycler v0.4.9b with default parameters. The average nucleotide identities were calculated from the assemblies using FastANI version 1.32 with default parameters. The reference genome of M. Abscessus strain ATCC 199977 was retrieved from NCBI under the accession number NZ_MLCG01000010.1.

*Mycobacterium tuberculosis mc*^*2*^*7000*, an attenuated strain of *Mycobacterium tuberculosis*, H37Rv, which carries deletions in RD1 and *panCD* loci, both of which are critical for virulence of *M. tuberculosis*, was grown at 37°C in Middlebrook 7H9 (DIFCO) supplemented with 10% OADC and 0.05% Tween 20 (Ojha et al., 2008).

### Antibiotic Sensitivity Assays

Wild type and mutant strains of *M. abscessus* were grown to an A_600_ of 0.6-0.7. Cells were tested for their susceptibility to rifampicin and rifabutin by spotting a 10-fold serial dilution on Middlebrook 7H10 (DIFCO) plates containing the indicated concentration of antibiotics. Antibiotic susceptibility in liquid media was assayed by inoculating the desired strain in a two-fold dilution series of each antibiotic at an initial A_600_ of 0.0004. The cultures were incubated at 37°C and the A_600_ was measured after 72 hours.

### RNA preparation / qPCR

*M. abscessus* (ATCC 19977) was grown to exponential phase (OD=0.4) in Middlebrook 7H9-OADC and exposed to a sublethal dose (16μg/ ml) of RIF for 30 mins. Total RNA was prepared using the Qiagen RNA preparation kit followed by DNAse I treatment. Unexposed cells were used as controls. Approximately 5 μg total RNA samples were treated with the Ribo-Zero™ rRNA removal procedure (Illumina) to enrich for mRNA. Approximately 500 ng of RNA was used for library preparation using the Script Seq v2 RNA-Seq kit and high throughput sequencing on the Illumina NextSeq platform. The sequence data was analyzed using the reference-based analysis and default parameters on Rockhopper v2.03 in which the data is normalized by upper quartile normalization and transcript abundance is reported as RPKM. Differential gene expression is tested for each transcript and q-values are then reported that control the false discovery rate (McClure et al., 2013; Tjaden, 2015). RNAseq experiments were performed three independent times, using two biological replicates each time.

*M. abscessus* strains (ATCC 19977, clinical strains and *ΔMab_helR*) were grown to A_600_ of 0.7, exposed to RIF(8μg/ml) or RBT (0.5 μg/ml) for 30mins. Total RNA was prepared using the Qiagen RNA preparation kit followed by DNAse I treatment. Primers for qRT-PCR were generated using Primer Quest software (IDT). cDNA was generated using random hexamers and Maxima reverse transcriptase (Fisher Scientific), and qRT-PCR performed using the Maxima SYBR Green qPCR Master Mix (Fisher Scientific) using the following primer pairs: *Mab_arr* 5’-CGTACTTCCATGGCAC CAA -3’/5’-GAATTTCTTGTCCGTCACGTTG-3’and *HelR* 5’-GGAGACGAACGTGGTGTTTA-3’/5’-TCGATCACAATGTGTCCATAGG. Applied Biosystems 7300 Real-Time PCR System was used with cycling conditions of: 50°C for 2 min, 95°C for 10 min, and 40 cycles of 95°C for 15 s, 60°C for 1 min. Data represents mean ± SD, n=3.

### Protein overexpression and purification

Wild type and mutant *helR* were cloned into pET21a with either a C-terminal his-tag or a C-terminal FLAG tag, transformed into BL21(DE3) pLysS, grown to an A_600_ of 0.5 and induced with 1mM IPTG at 30°C. The cells were lysed in a buffer containing 50mM Tris-HCl (pH 8.0), 300mM NaCl, 1mM MgCl_2_, 5% glycerol, 1mM PMSF and 0.25% sodium deoxycholate and the clarified lysate was loaded on a Ni-NTA column (Qiagen). Non-specifically bound proteins were removed by washing with lysis buffer containing 20mM imidazole and the proteins eluted with buffer containing 150mM imidazole. For purification of FLAG-tagged proteins, cells were lysed in a buffer containing 50mM Tris-HCl (pH 8.0), 150 mM NaCl, 1mM MgCl_2_, 5% glycerol and mixed with Anti-FLAG M2 magnetic beads (Sigma). Following wash with the same buffer, FLAG-tagged proteins were eluted by competition using the FLAG peptide (Sigma).

*M. tuberculosis* σ^A^ cloned in pET21a with a C-terminal his –tag was transformed into BL21(DE3) pLysS, grown to an A_600_ of 0.4 and induced with 1mM IPTG at 30°C. The cells were lysed in a buffer containing 50mM Tris-HCl (pH 8.0), 300mM NaCl and 5% glycerol and the clarified lysate was loaded on a Ni-NTA column (Qiagen). Non-specifically bound proteins were removed by washing with lysis buffer containing 40mM imidazole and σ^A^ eluted with 150mM imidazole. For purification of *M. tuberculosis* RNA polymerase, BL21(DE3) pLysS was co-transformed with pETDuet-Mtbββ’ and pRsfDuet-Mtbαω, grown at 30°C to an A_600_ of 0.4 and induced with 0.4mM IPTG at 16°C for a period of 18hours. The cells were lysed by sonication and passed through a Ni-NTA column (Qiagen) equilibrated with 50mM Tris, 300mM NaCl, 1mM MgCl_2_, and 5% glycerol, protease inhibitor cocktail (Lysis buffer). The column was washed Lysis Buffer + 40mM imidazole and eluted with Lysis Buffer +150mM imidazole. Fractions containing RNAP were loaded on a Heparin Sepharose matrix (GE Healthcare) equilibrated with 50mM Tris, 300mM NaCl and 5% glycerol and eluted with a buffer containing 1M NaCl.

For purification of *M. abscessus* RNA polymerase, MabATCC-rpoA_his_ in which the native *rpoA* was tagged at the C-terminal with 10X-his was grown to an A_600_= 0.8. Cells were treated with 16μg/mL RIF for 30 mins when required. The cells were harvested and lysed using The CryoMill (Retsch) in a buffer containing 50mM Tris, 200mM NaCl, 5% glycerol, 1mM MgCl_2_, protease inhibitor cocktail and loaded onto a Ni-NTA matrix equilibrated with the above buffer containing 5mM imidazole. Non-specific proteins were removed by washing with buffer containing 40mM imidazole followed by elution with 150mM imidazole buffer.

### Immunoprecipitation and Western Blotting

MabATCC-*helR*_FLAG_ in which the native *helR* was tagged at the C-terminal with 3X-FLAG was grown to an A_600_= 0.8. Cultures were treated with 8-16 μg/mL RIF for 30 mins and cells were harvested and lysed using The CryoMill (Retsch) in a buffer containing 50mM Tris, 50mM NaCl, 5% glycerol, 1mM MgCl_2_, protease inhibitor. The lysate was clarified by centrifugation, protein concentration was determined at A_260_ and equal quantities of protein from different samples were mixed with Anti-FLAG M2 magnetic beads (Sigma), washed with 20 bed volumes to remove non-specifically bound proteins and eluted using 0.1M glycine, pH 3.0. The eluates were neutralized using 1M Tris, pH 8.0, separated using 8% SDS-PAGE transferred onto a PVDF membrane and probed with anti-FLAG (Sigma) and anti-RpoA (Biolegend) monoclonal antibodies. Purified Mtb RNAP was used as a control.

### *In vitro* transcription Assays

Multiple round *in vitro* transcription was performed as previously described (Huang et al., 2012). In short, 75nM *M. tuberculosis* RNAP was assembled with 300nM purified σ^A^ in a volume of 10μl for 10 minutes at 37°C. RIF was added to indicated concentrations for 15 mins at 37°C. 25nM of *sinP3* promoter DNA was added to the mixtures and incubated for 10 mins at 37°C. Purified MabHelR was added as indicated. Transcription was initiated by addition of 2μL of NTP mix (1.5mM of ATP, GTP and CTP and 0.5 mM UTP) + 2μCi of ^32^P-α-UTP. Reactions were incubated at 37°C for 30 mins and terminated by the addition of 5mM EDTA + 100μg/ml tRNA. Samples were ethanol precipitated, resuspended in STOP buffer (80% v/v Formamide/10mM EDTA/0.01% Xylene Cyanol/0.01% Bromophenol Blue) and separated using denaturing PAGE (6% Urea polyacrylamide gel). The products were visualized using a Typhoon Imager (GE Healthcare) and quantitated using the Image Quant software.

### Generation of RIF-stalled RNAP complexes and disassembly

RNAP stalled in the presence of RIF was generated by using a modification of a previous protocol for trapping stalled elongation complexes (Kouba et al., 2020). First, holoenzyme was formation was initiated by mixing 32 pmoles of MabRNAP with a 4-fold molar excess of MabSigA and incubation at 37°C for 5mins. The samples were mixed with 8 pmoles of double-stranded *sinP3* DNA that was biotinylated at the 5’ end in buffer containing 40mM Tris, pH 8.0, 10mM MgCl_2_ and 1mM DTT for 5 mins followed by addition of 90pmoles of RIF and 1mM ATP for 10 mins at 25°C to enable formation of stalled RNAP complexes. Streptavidin coated magnetic beads (Pierce, 25 μl per sample) was washed with 500μl of binding buffer (20mM Tris, pH 8.0, 100 mM NaCl), resuspended in 25 μl of the same buffer. The assembled stalled complexes were mixed with washed buffer and incubated at 25°C for 20 mins. The beads were washed with 500μl wash buffer (20mM Tris, pH 8.0, 100 mM NaCl, 2mM MgCl_2_ and 1mM DTT) followed by a wash with 500μl reaction buffer (40mM Tris, pH 8.0, 10mM MgCl_2_, 100mM KCl and 1mM DTT). The beads were resuspended in 10μl reaction buffer containing WT or mPCh mutant of HelR with or without ATP (1mM) and incubated at 37°C for 30 mins. The eluted samples were separated on an 8% SDS-PAGE, transferred to a PVDF membrane and probed with anti-his antibodies that recognizes the his-tagged RpoA. Three biological replicates off the experiment was conducted and the western blots were quantitated using ImageJ (Schneider et al., 2012).

### ATP hydrolysis assay

ATPase activity of Wt and mutant HelR proteins was determined using the colorimetric ATPase assay kit (Sigma). The ATP hydrolysis reaction was performed by incubating 0.3 µM HelR with 1 mM ATP (NEB), in a buffer containing 40 mM Tris, 80 mM NaCl, 8 mM MgAc2, 1 mM EDTA, pH 7.5 for 30mins at room temperature followed by addition of the malachite green reagent included in the kit. Following incubation for 30 mins at room temperature to generate the colorimetric product, the absorbance of the samples was measured at 620nm. The amount of free phosphate liberated was determined from a standard curve that was generated using phosphate standards provided in the kit.

### Protein identification through mass spectrometry

LC MS/MS was performed by the RNA Epitranscriptomics & Proteomics Resource at the University at Albany. Briefly, the protein band corresponding in size to MabHelR in RNAP preparations was manually excised, minced, in-gel digested with trypsin (Sigma, St. Louis, MO) and extracted three times with 50% acetonitrile containing 5% formic acid. LC-MS/MS was performed on an integrated micro LC-Orbitrap Velos system (Thermo). Tandem spectrum data was processed using Mascot 2.7 (Matrix Science, Boston, MA). The list of peaks was used to query the Mycobacteriaceae protein database downloaded from NCBI by setting the following parameters: peptide mass tolerance, 10 ppm; MS/MS ion mass tolerance, 0.1 Da; allowing up to one missed cleavage; considering variable modifications such as methionine oxidation, cysteine carboxyamidomethylation, and deamidation. Only significant hits as defined by Mascot probability analysis was considered for a positive protein identification.

## Acknowledgements

We thank The Wadsworth Center’s Applied Genomics Technology Core, the Bioinformatics Core and the Media Core. We thank Qishan Lin from The University at Albany for performing LC MS/MS for protein identification. We also thank Jon Paczkowski and Anil Ojha for invaluable discussions and critical reading of the manuscript. PG is supported by NIH awards AI155473 and AI146774 and the Wadsworth Center.

## Supplementary Figures

**Figure S1:**
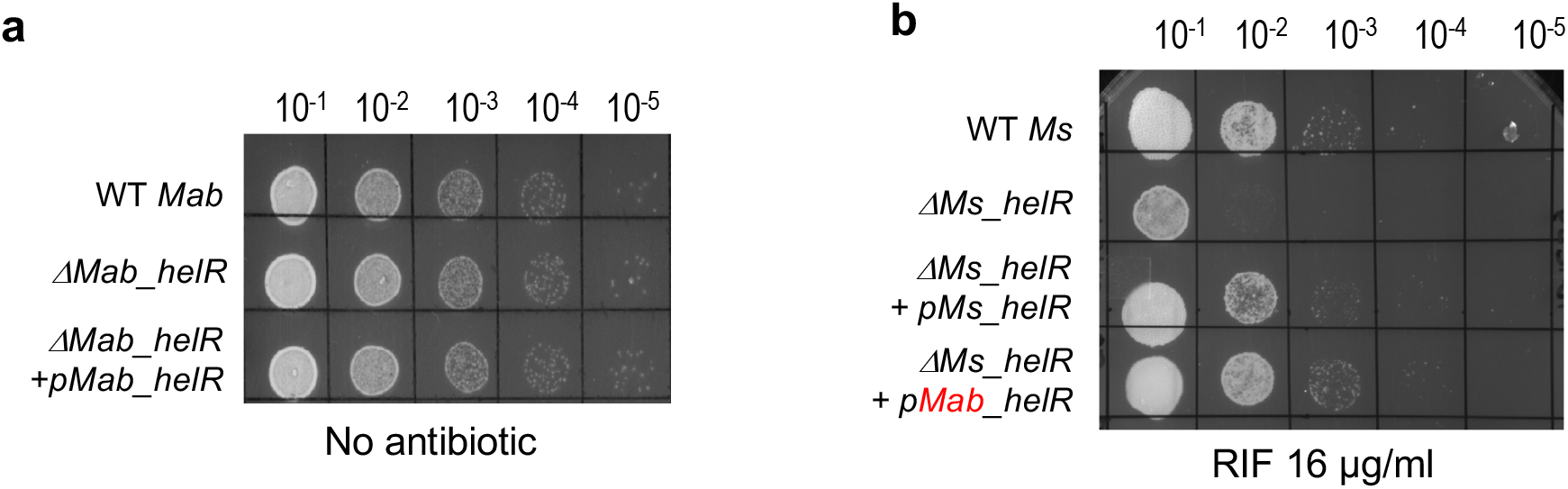
**a)**Wild type *M. abscessus, ΔhelR* and complemented strains were grown to A_600_ of 0.7 and a 10-fold dilution series was spotted on media lacking antibiotics. No growth defect was observed in the *ΔhelR* mutant compared to WT bacteria. **b)** Wild type *M. smegmatis, ΔMs_helR* and *ΔMs_helR* complemented with either *HelR* or *Ms_helR* were grown to A_600_ of 0.7 and a 10-fold dilution series was spotted on media RIF (16μg/mL). Mab_helR can complement the RIF sensitivity of *ΔMs_helR*.

**Figure S2:**
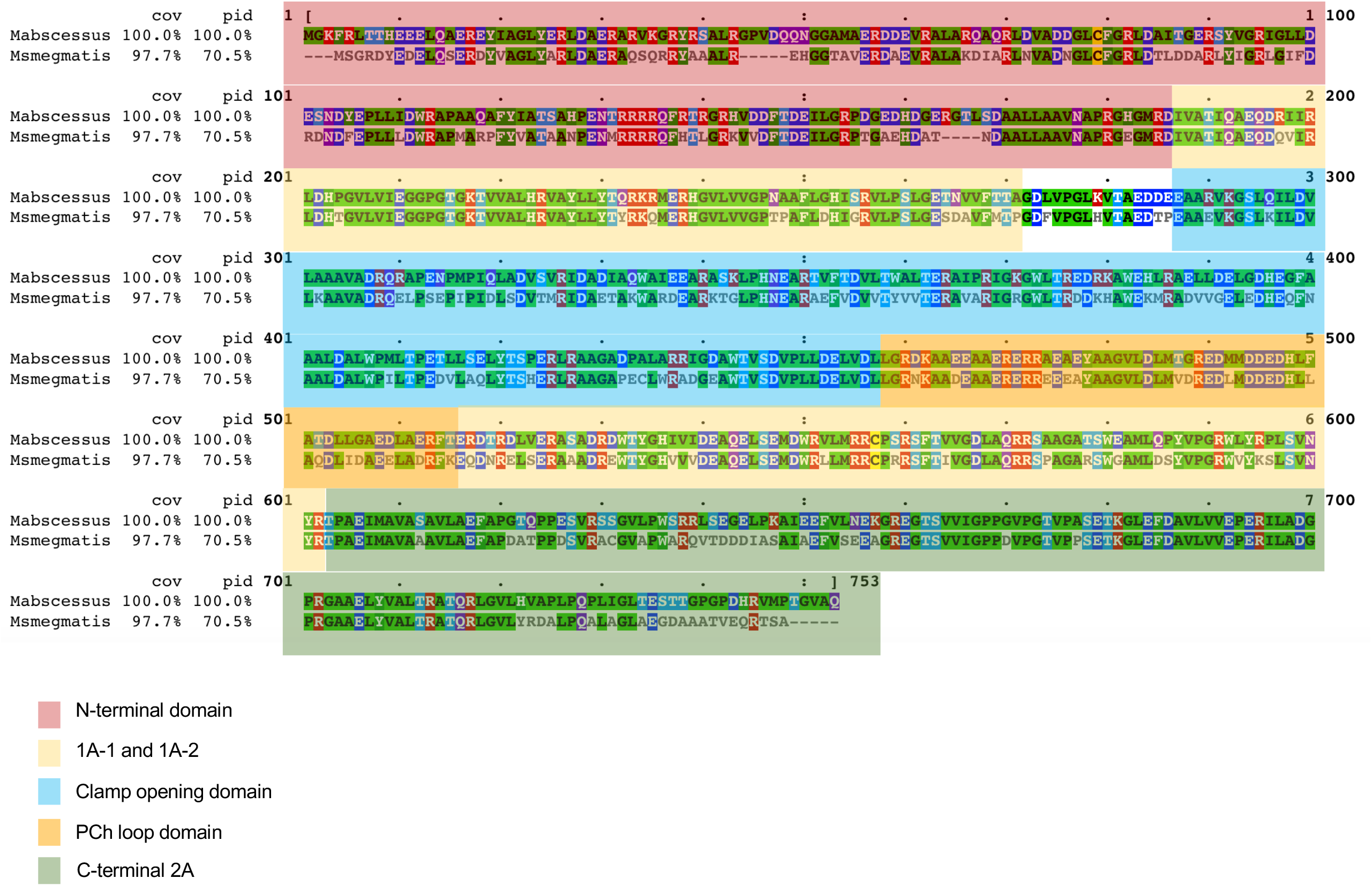
Sequence comparison of HelR from *M. abscessus and M. smegmatis*. Sequence alignment was performed using CLUSTALW. The N-terminal, C-terminal, 1A-1 and 1A-2 and the HelR specific domains are shaded.

**Figure S3:**
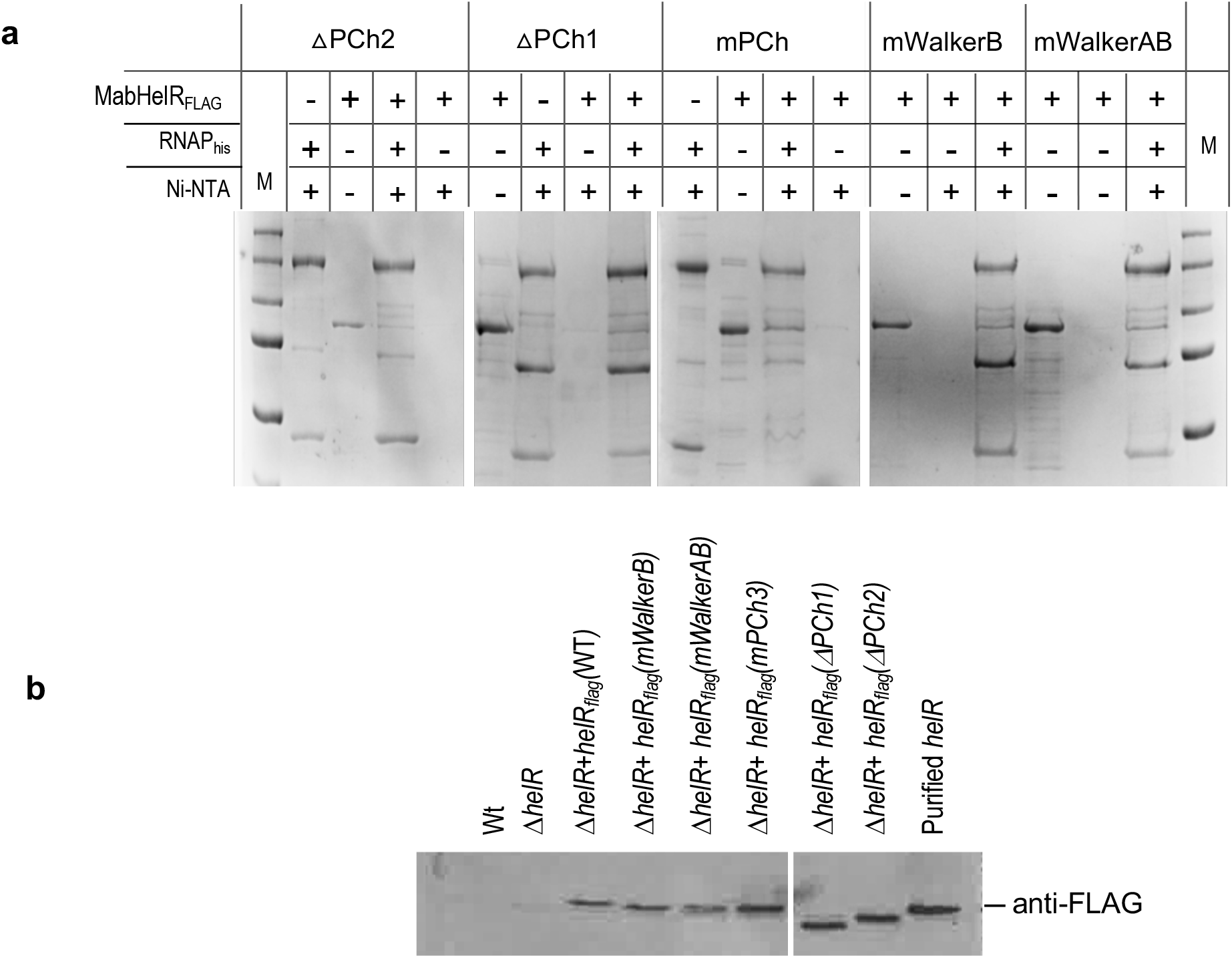
Expression of mutant HelR *in vivo* and interaction with RNAP *in vitro*. **b)** Immunoblot with anti-FLAG antibodies of *M. abscessus* ATCC 19977, *ΔMab_helR* and *ΔMab_helR* complemented with either *WT helR*_*FLAG*_, *helR* (ΔPCh1) _*FLAG*_, *helR* (ΔPCh2) _*FLAG*_ and *helR* (mPCh) _*FLAG*_ treated with RIF (16μg/mL) shows that WT and mutant proteins are induced to similar levels in the complementing strains. **c)** Coomassie stained gel showing elution of proteins using nickel affinity chromatography. Interaction assays were carried out using WT and mutant HelR_FLAG_ with RNAP_his_ as indicated and followed by binding and elution from a Ni-NTA column. Control samples included either only RNAP_his_ or HelR_FLAG_. Purified RNAP_his_ or HelR_FLAG_ were included as markers.

**Table S1.** MIC of RIF and RBT of *M. abscessus* clinical strains.

| Strain | Subspecies | Colony Morphology | Minimum Inhibitory Conc. (µg/mL) |  |
| --- | --- | --- | --- | --- |
|  |  |  | RIFAMPICIN | RIFABUTIN |
| ATCC19977 | *M. abscessus abscessus* | smooth | 64 | 2 |
| Mab 095 | *M. abscessus massiliense* | smooth | 64 | 2 |
| Mab 174 | *M. abscessus massiliense* | smooth | 64 | 2 |
| Mab 299 | *M. abscessus abscessus* | rough | 64 | 2 |
| Mab 427 | *M. abscessus abscessus* | n/a | 32 | 1 |
| Mab 525 | *M. abscessus abscessus* | rough | 128 | 4 |
| Mab 549 | *M. abscessus bolletii* | smooth | >64 | 4 |
| Mab 698 | *M. abscessus massiliense* | smooth | 64 | 2 |
| Mab 998 | *M. abscessus abscessus* | smooth | 32 | 1 |

**Table S2.**
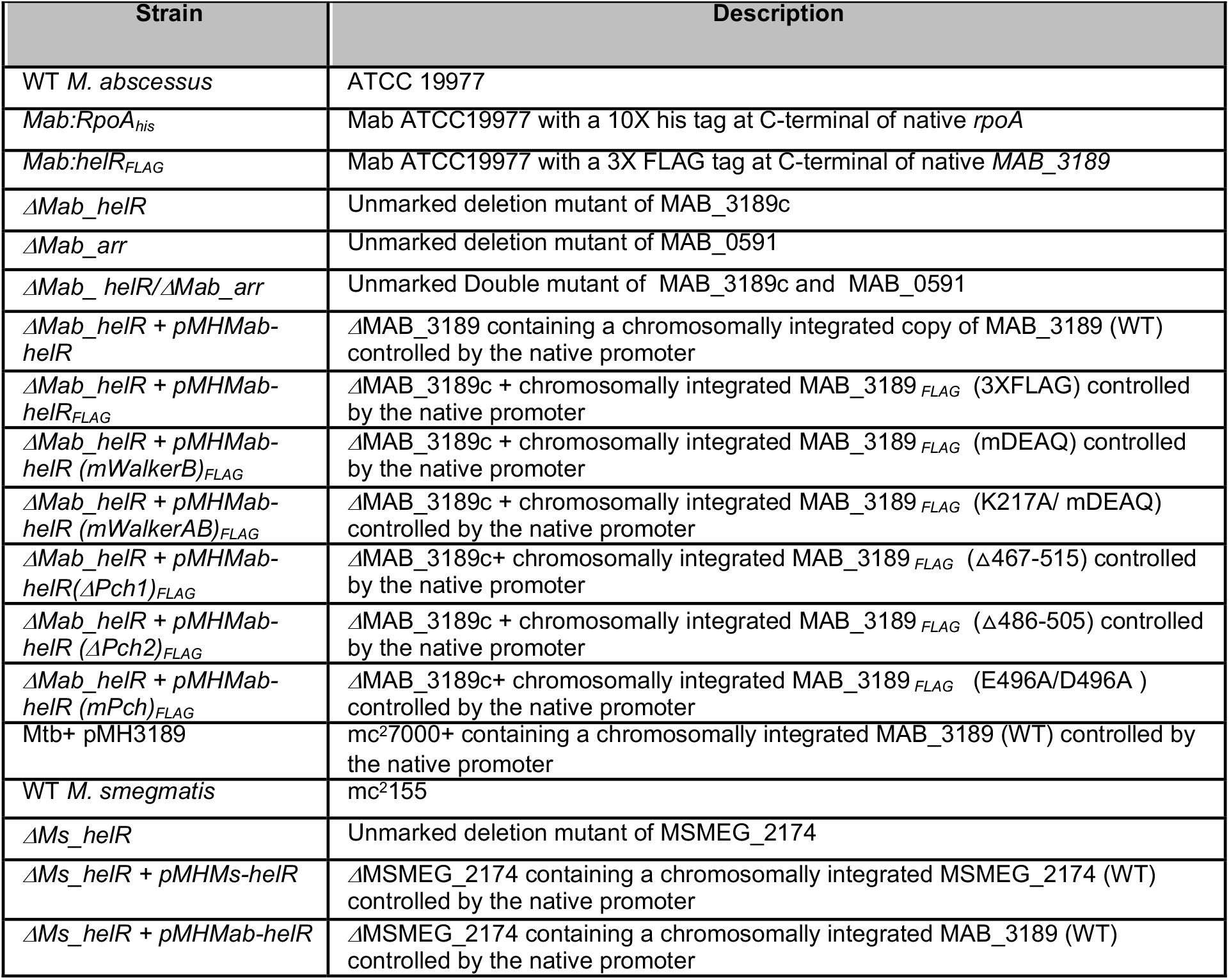
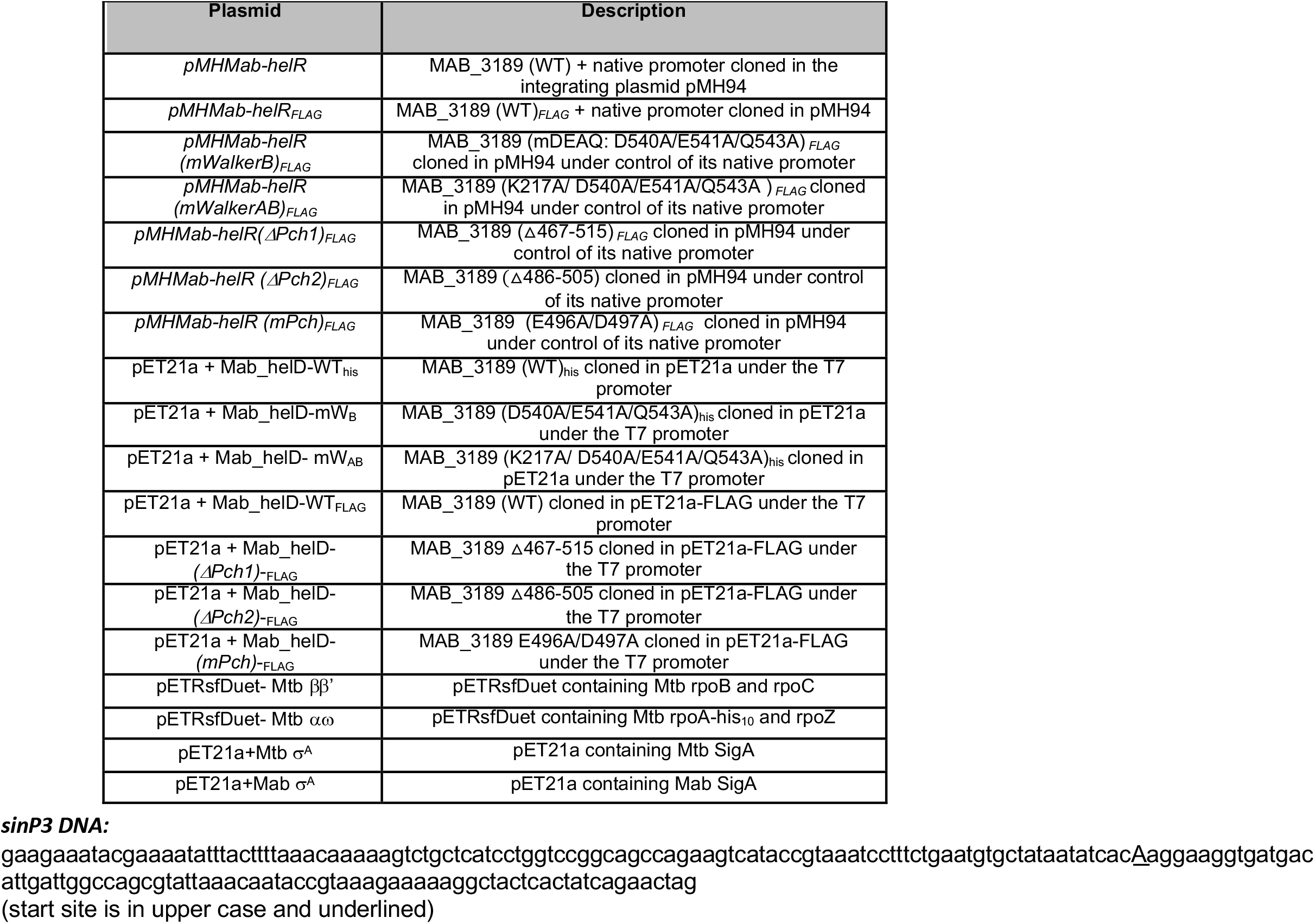
List of strains and plasmids used in the study.

